# Clinical and biomarker changes in sporadic Alzheimer’s disease

**DOI:** 10.1101/2020.05.08.084293

**Authors:** Junjie Zhuo, Yuanchao Zhang, Bing Liu, Yong Liu, Xiaoqing Zhou, Perry F. Bartlett, Tianzi Jiang, the Alzheimer’s Disease Neuroimaging Initiative

**Affiliations:** Brainnetome Center, Institute of Automation, Chinese Academy of Sciences, Beijing 100190, China; The Queensland Brain Institute, University of Queensland, Brisbane, QLD 4072, Australia; University of Chinese Academy of Sciences, Beijing 100049, China; CAS Center for Excellence in Brain Science, Institute of Automation, Chinese Academy of Sciences, Beijing 100190, China; The Clinical Hospital of Chengdu Brain Science Institute, MOE Key Lab for Neuroinformation, University of Electronic Science and Technology of China, Chengdu 625014, China

## Abstract

**IMPORTANCE:** The dynamic changes of biomarkers and clinical profiles in sporadic Alzheimer’s disease (SAD) are poorly understood.

**OBJECTIVE:** To evaluate the impact of amyloid-β (Aβ) biomarkers on SAD by measuring the dynamic changes in biomarkers and clinical profiles in the progression of SAD.

**DESIGN AND SETTING:** This retrospective and longitudinal study analyzed clinical and biomarker data from 665 participants (mean follow-up 4.90 ± 2.83 years) from a subset of the AD Neuroimaging Initiative (ADNI) participants collected from August 2005 to December 2018. By aligning the timing of the changes in the various biomarkers with the stable normal cognition (CN) baseline and mild cognitive impairment (MCI) or AD onset timepoints, we combined data from the stable CN, CN conversion to MCI (CN2MCI), and MCI conversion to AD (MCI2AD) groups to identify the trajectories associated with the progression of AD.

**PARTICIPANTS:** The participants were 294 CN, 69 CN2MCI, 300 MCI2AD, and 24 who converted from CN to MCI to AD (CN2MCI2AD) (of whom 22 were also included in the CN2MCI).

**EXPOSURES:** Amyloid-β measured by florbetapir positron emission tomography (PET) or cerebrospinal fluid assay of amyloid-β (CSF Aβ_42_).

**MAIN OUTCOMES AND MEASURES:** The measures included the 13-item cognitive subscale of the AD Assessment Scale (ADAS13, as a clinical measure), hippocampal volume, and the fluorodeoxyglucose (FDG) PET standardized uptake value ratio (SUVR).

**RESULTS:** The CN, CN2MCI, and MCI2AD subgroups’ median (interquartile range [IQR]) annual changes in ADAS13 were (0.388 [−0.278, 0.818], 1.000 [0.239, 2.330], and 3.388 [1.750, 6.169]). The annual changes in hippocampal volume for each group were (−0.005 %ICV [−0.011, −0.001], −0.006 %ICV [−0.012, −0.002], and −0.014 %ICV [−0.021, −0.009]). The annual changes in FDG PET SUVR for each group were (−0.011 [−0.030, 0.010], −0.027 [−0.056, −0.012], and −0.039 [−0.063, 0.014]). Changes in the amyloid biomarkers were inconsistent with clinical profile changes. The annual changes in CSF Aβ_42_ for each group were (−1.500 pg/ml [−6.000, 4.000], −2.200 [−5.667, 4.000], and −2.000 [−7.000, 2.650]) and in Aβ PET SUVR for each group were (0.004 [−0.002, 0.012], 0.004 [−0.001,0.011], and 0.005 [−0.006, 0.014]). During the stable CN and CN2MCI stages, subjects with elevated and those with normal amyloid showed no significant differences (likelihood ratio test, *p* < .01) in clinical measures, hippocampal volume, or FDG.

**CONCLUSIONS AND RELEVANCE:** Hippocampal volume and FDG associated with clinical profiles impairment in the SAD progression. Aβ alone is not associated with clinical profiles, hippocampal volume, and FDG impairment in the preclinical stage of SAD.

**Key Points:** **Question:** What is the role of amyloid-β in dynamic changes in biomarkers and clinical profiles in the progression of sporadic Alzheimer’s disease?

**Findings:** The changes of the hippocampal volume and FDG that were consistent with the changes of the clinical profiles showed a non-linear change in the initial stage and an accelerated non-linear change during MCI2AD, changes in amyloid biomarkers were inconsistent with the clinical profile. Cognitively normal people with elevated or normal amyloid showed no significant differences in clinical measures, hippocampal volume, or FDG.

**Meaning:** Amyloid-β alone may not be used as the central index for defining the preclinical stage of SAD.

## Introduction

The diagnostic guidelines for Alzheimer’s disease (AD)^1,2^ provide a clinical-pathological framework. The National Institute on Aging-Alzheimer’s Association (NIA-AA), in line with the amyloid hypothesis^3,4^, defines AD on the basis of biomarkers, rather than by clinical symptoms^5^. However, two observations, the failure of all anti-amyloid-β (Aβ) drugs^6–9^ to show clinical efficacy and the discovery that amyloid plaques are not unique to AD^5,10^, have led to a debate about the central role of amyloid in the etiology of the disease and its usefulness as a diagnostic marker of AD.

To address this debate, identifying which dynamic changes in biomarkers and clinical profiles correlate directly with the progression of AD is essential. The relevant studies have primarily relied on patients with autosomal dominant AD (ADAD), who often have a predictable age at onset^11–13^. In contrast, the precise timing of the disease for patients with sporadic AD (SAD) is difficult to predict^14^. Because the ADAD genetic mutations (*APP, PSEN1,* and *PSEN2*) cause alterations in Aβ processing, ADAD studies have consistently found that Aβ is the first and key biomarker, followed by changes in other biomarkers and clinical profiles^11–13^. However, increasing evidence has shown that patients with SAD are associated with multiple gene factors, which affect more than Aβ processing^10,15–17.^ Since ADAD only accounts for a very small proportion (approximately 1%) of AD^11^, how widely applicable the findings obtained from ADAD to SAD remains a question^10^.

A previous prospective SAD study based the stage of AD on the level of accumulation of amyloid and found, consistent with ADAD studies, that the Aβ abnormality appeared first, followed by other changes^18^. However, due to its assumption that Aβ is the etiological agent, that study does not consider the possible dynamic biomarker and clinical changes which occur in relation to symptom onset as in the previous ADAD studies^11–13^. Even in subjects who have over 15 years of longitudinal data, the baseline has not been aligned with the onset of clinical symptom to investigate longitudinal changes in biomarkers and clinical profiles^19^. However, as the progression of AD has been hypothesized to be non-linear^20,21^, simply aligning the baseline with Aβ levels or studying the longitude data is not sufficient to chart the progression of SAD. Thus, to reduce this limitation, we aligned the timepoints of the clinical diagnosis of mild cognitive impairment (MCI) or AD onset to investigate the dynamic changes that occur from cognitively normal (CN) to MCI and from MCI to AD.

## Methods

### Study design

The data were obtained from the ADNI dataset (http://adni.loni.usc.edu/) and downloaded in December 2018. The ADNI was launched in 2003 as a public-private partnership, led by Principal Investigator Michael W. Weiner, MD. The primary goal of ADNI has been to test whether serial magnetic resonance imaging (MRI), positron emission tomography (PET), other biological markers, and clinical and neuropsychological assessment can be combined to measure the progression of MCI and early AD.

To estimate the timing, order, and trajectory of clinical and biomarker changes from normal aging to AD, we labeled the data of the three subgroups as CN, subjects with normal cognition who were confirmed to convert to MCI (CN2MCI), and subjects with MCI who were confirmed to convert to AD (MCI2AD). The CN subgroup was defined as either subjects who had a cognitively normal baseline, showed no significant memory concern (SMC), and had at least two years’ follow-up without conversion to MCI or AD or as subjects with a baseline of MCI who reversed to CN within one year and remained stable CN for at least 2 years to the end of follow-up. The CN2MCI subgroup was defined as subjects with a baseline diagnosis of cognitively normal and a subsequent diagnosis of having converted to MCI in the follow-up or as subjects with a SMC confirmed as having converted to MCI. To increase the sample size and statistical power, the CN2MCI timepoint of subjects who converted to MCI and finally to AD were also included in the CN2MCI group. The MCI2AD subgroup was defined as subjects with a baseline diagnosis of MCI who converted to stable AD in the follow-up.

To precisely reflect the stage of disease, we selected those subjects within the CN2MCI and MCI2AD subgroups who had one year or less between the initial one-time assessment before the disease onset and the disease onset of MCI or AD.

### Assessments

The clinical profiles and biomarkers used in the present study included the 13-item cognitive subscale of the Alzheimer’s Disease Assessment Scale (ADAS13), Mini-Mental State Examination (MMSE), Clinical Dementia Rating Scale-Sum of Boxes (CDRSB), hippocampal volumes, fluorodeoxyglucose (FDG) positron emission tomography (PET), florbetapir PET, and CSF biomarkers (including tau, phosphor-tau (Ptau), and Aβ_42_). FDG and florbetapir PET, were measured by the standardized uptake value ratio (SUVR). Please see the Supplementary materials for all the metadata downloaded from the ADNI dataset and the detailed assessment of each clinical profile and biomarker.

Participants were categorized into elevated amyloid or normal amyloid subsets depending on their florbetapir SUVR or CSF Aβ_42_ status. Elevated amyloid was defined as a florbetapir SUVR greater than 0.79^22^ or a CSF Aβ_42_ value less than 192 pg/mL^23^. Participants were classified as having elevated amyloid if they met the cutoff threshold at any timepoint. Otherwise, they were classified as having normal amyloid. If there was no amyloid information for a participant, their data were classified as missing.

As the ADAS has usually been used to monitor the progression of AD^8,24^, we calculated the correlations between the ADAS13 and each marker in the CN2MCI and MCI2AD stages separately to evaluate whether the markers could predict AD progression.

To compare the progression curve for all the markers and verify the model of the fitted results, the scaled value for each marker was defined by (raw data – mean CN baseline value) / the standard deviation (SD) of the whole dataset. To further verify the abnormal pattern of the markers in the progression of AD, we also analyzed the within-individual trajectories for all 24 subjects who were initially diagnosed as CN, subsequently converted to MCI, and then to AD (CN2MCI2AD). Each marker in these individuals was also scaled by the mean of the baseline data for the CN subgroup and for the SD of the entire dataset.

### Statistical analysis

For the longitudinal trajectory analyses of the CN2MCI and MCI2AD subgroups, the follow-up years were categorized into pre-symptom onset (<0 onset years) and post-symptom onset (>0 onset years). To increase model convergence, we excluded the data of timepoints for which the sample size was less than 3 for each clinical profile or biomarker, (See Fig. S1 for the detailed sample size for the various timepoints for each clinical profile or biomarker) Statistical analyses and plotting were performed using R (version 3.5.3, https://www.r-project.org/)

Longitudinal trajectory models were constructed for the various biomarkers using linear mixed effects models^25^. For each marker, we started by fitting an appropriate function to the time (baseline or onset time) e.g. time + time^^2^ + time^^3^. Disease progression (CN, CN2MCI, and MCI2AD) was included in the models to extract disease-specific biomarker trajectories. Covariates such as age at baseline or onset year, sex, APOEε4, and education were included as confounds, and a backward elimination method was used for model selection. We then selected a structure for the random effects and covariance structure for the residuals in the model. All the model selections were based on the Akaike Information Criterion^14,26^, an objective model selection tool. Maximum likelihood was used to fit the mixed-effect models as it is robust to the absence of random data^27^.

We further compared the trajectories for each marker in the progression of AD to uncover differences between the elevated amyloid and normal amyloid groups. The overall amyloid effect was tested using likelihood ratio tests that compared the full model to a reduced model with no amyloid factor in each subgroup for each marker. For any subgroup that showed a significant amyloid effect as the disease progressed, a supplementary *post hoc* analysis was performed between the elevated amyloid and normal amyloid groups at each timepoint based on the estimated marginal means derived from the model.

To determine the timing of the dysfunctions, we fitted a linear mixed effects model to the CN2MCI subgroup with time as a categorical variable for each biomarker. The *post hoc* analysis was conducted between each timepoint based on estimated marginal means derived from the model.

## Results

Of the downloaded 665 subjects from the ADNI dataset, we utilized the data from 663 participants in the group analysis (CN: 294, CN2MCI: 69, and MCI2AD: 300, for more details of the participants’ characteristics see Table 1) and 24 CN2MCI2AD participants in the individual analysis (the data from the group of 22 participants in the CN2MCI stage was combined into the above CN2MCI group analysis). Some participants were followed for up to 13 years with a mean follow-up period of 4.90 ± 2.83 years.

**Table 1.** Characteristics of Study Participants Abbreviations: ADASA13, the 13-item cognitive subscale of the Alzheimer’s Disease Assessment Scale; CDRSB, the Clinical Dementia Rating Scale-Sum of Boxes; MMSE, the Mini-Mental State Examination; ICV, intracranial volume; FDG, fluorodeoxyglucose; CSF, cerebrospinal fluid; SUVR, standardized uptake value ratio; SD, standard deviation

|  | CN | CN2MCI | MCI2AD | *p* |
| --- | --- | --- | --- | --- |
|  | N=294 | N=69 (22 finally to AD) | N=300 |  |
| Sex: |  |  |  | .058 |
| Female | 146 (49.7%) | 28 (40.6%) | 121 (40.3%) |  |
| Male | 148 (50.3%) | 41 (59.4%) | 179 (59.7%) |  |
| Education, mean (SD), y | 16.5 (2.68) | 16.1 (2.67) | 15.9 (2.80) | .032 |
| APOE allele: |  |  |  | <.001 |
| APOEε4 noncarriers | 221 (75.2%) | 42 (60.9%) | 98 (32.7%) |  |
| APOEε4 carriers | 73 (24.8%) | 27 (39.1%) | 202 (67.3%) |  |
| Follow-up, mean (SD), y | 5.33 (2.83) | 5.70 (3.43) | 4.07 (2.29) | <.001 |
| Amyloid characteristics: |  |  |  | <.001 |
| Missing amyloid information | 52 (17.7%) | 8 (11.6%) | 75 (25.0%) |  |
| Elevated amyloid | 115 (39.1%) | 41 (59.4%) | 202 (67.3%) |  |
| Normal Amyloid | 127 (43.2%) | 20 (29.0%) | 23 (7.7%) |  |
|  | Baseline characteristics | MCI onset characteristics | AD onset characteristics |  |
| Age, mean (SD), y | 74.0 (6.18) | 79.7 (5.70) | 76.5 (7.39) | <.001 |
| ADAS13, mean (SD) | 8.60 (4.04) | 15.0 (6.32) | 27.1 (7.36) | <.001 |
| CDRSB, mean (SD) | 0.06 (0.26) | 1.01 (0.76) | 4.30 (1.59) | <.001 |
| MMSE, mean (SD) | 29.1 (1.17) | 27.6 (1.83) | 23.8 (2.91) | <.001 |
| Hippocampal volume, mean (SD), %ICV | 0.50 (0.06) | 0.44 (0.05) | 0.37 (0.06) | <.001 |
| FDG PET SUVR, mean (SD) | 1.41 (0.14) | 1.20 (0.15) | 1.16 (0.14) | <.001 |
| Amyloid PET SUVR, mean (SD) | 0.78 (0.09) | 0.93 (0.14) | 1.01 (0.12) | <.001 |
| CSF Aβ42, mean (SD), pg/ml | 207 (49.9) | 194 (79.0) | 137 (35.6) | <.001 |
| CSF tau, mean (SD), pg/ml | 63.8 (29.2) | 90.4 (28.8) | 130 (75.9) | <.001 |
| CSF ptau, mean (SD), pg/ml | 29.1 (14.7) | 39.2 (23.7) | 55.1 (30.7) | <.001 |

Figure 1 shows the trajectories of the biomarkers estimated by the linear mixed effects models across groups (for the spaghetti plot of the raw data, see eFigure 2). Consistent with the clinical profiles of AD progression, the hippocampal volume and FDG levels remained stable throughout the CN stage followed by slow, non-linear changes in the CN2MCI stage and rapid non-linear changes in the MCI2AD. Even the values of the florbetapir PET and the CSF biomarkers were normal in CN, intermediate in CN2MCI and abnormal in MCI2AD subgroups, the shape of the trajectories for florbetapir PET and the CSF biomarkers did not consistent with the clinical profile. The details of the linear mixed model for each biomarker are displayed in eTables 1-9.

**Figure 1.**
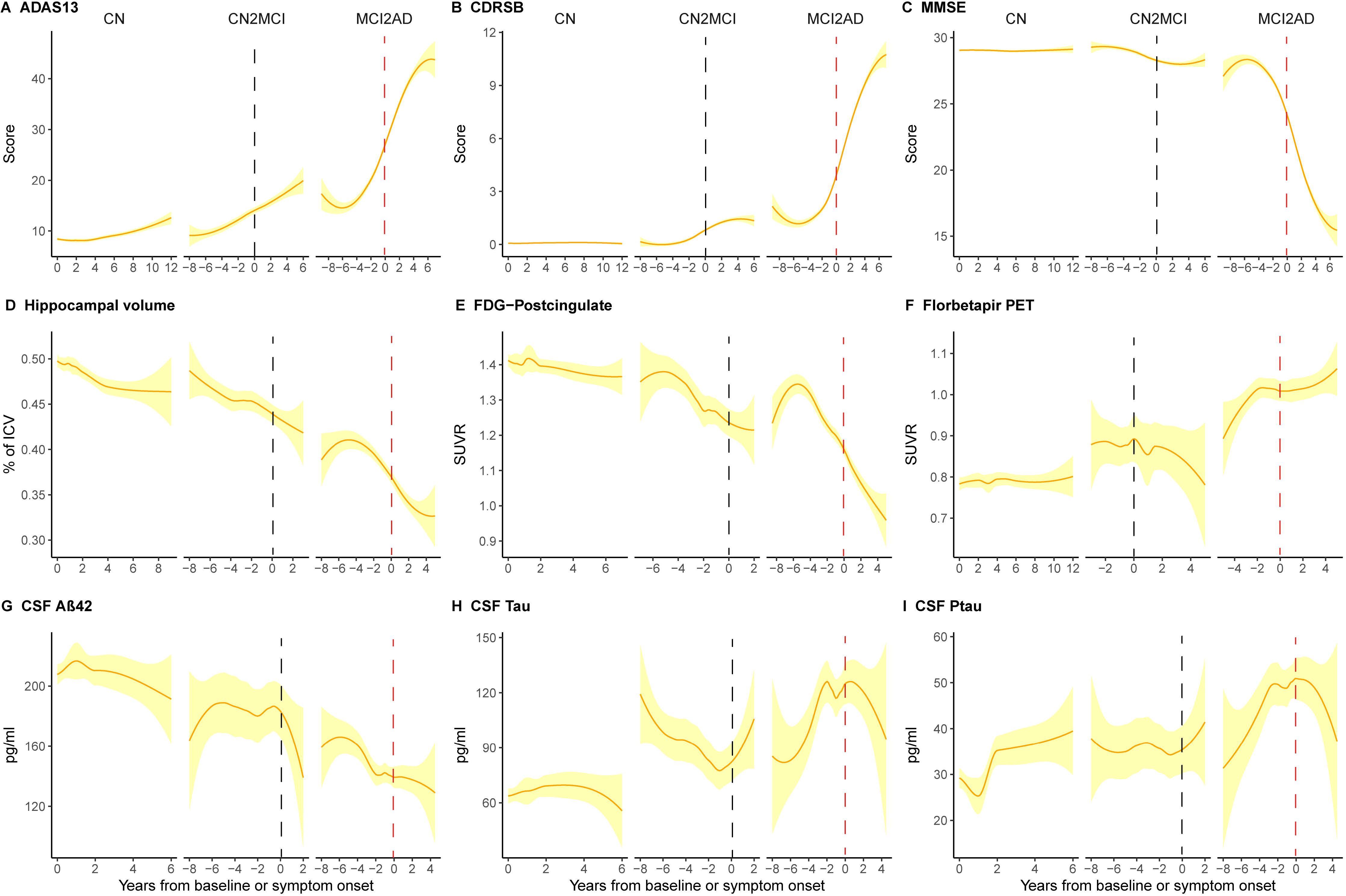
Estimated group trajectories of clinical profiles and biomarkers. (A) ADAS13; range from 0 [best] to 85 [worst], (B) CDRSB; range from 0 [best] to 18 [worst], (C) MMSE; range from 0 [worst] to 30 [best], (D) the MRI measures of hippocampal volumes adjusted by percent of the total intracranial volume (ICV), (E) the post-cingulate cortex glucose metabolism measured by fluorodeoxyglucose (FDG) positron emission tomography (PET) consistently showed stable changes in the stable cognitive normal (CN) subgroup, slow non-linear changes in the confirmed CN conversion to MCI (CN2MCI) subgroup, and acceleration non-linear changes in the confirmed MCI conversion to AD (MCI2AD) subgroup. In contrast, (F) Florbetapir PET, (G) CSF Aβ_42_, (H) CSF tau, and (I) CSF phosphor-tau (Ptau) did not show changes consistent with the clinical profiles. The estimated trajectory and 95% confidence interval from the linear mixed models (yellow line and yellow shaded area, respectively) are plotted against years from baseline or symptom (MCI or AD) onset for each marker. The black dashed line represents the MCI onset timepoint. The red dashed line represents the AD onset timepoint.

**Figure 2.**
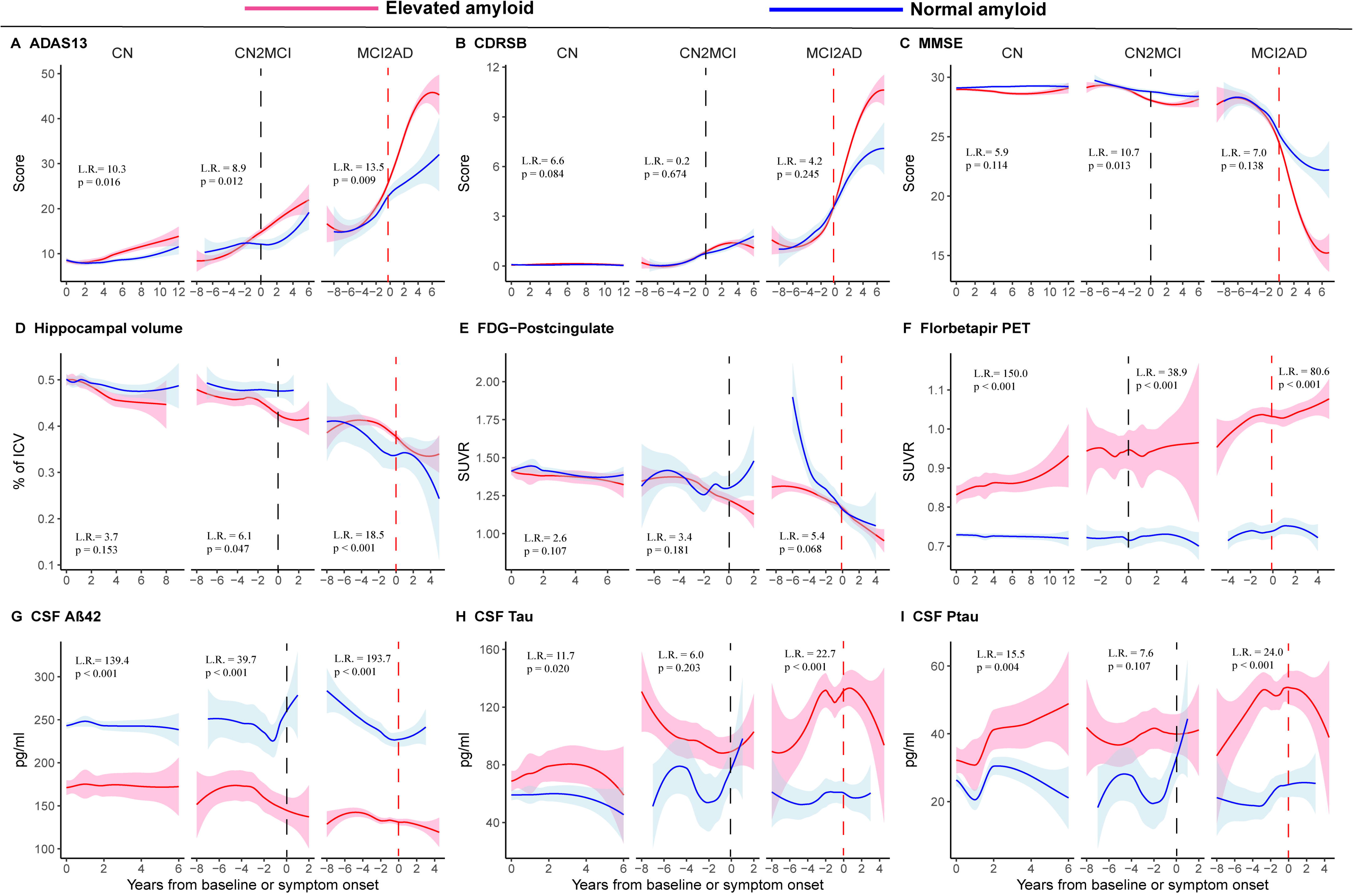
Estimated elevated and normal amyloid group trajectories of clinical profiles and biomarkers. See Figure 1 for explanation of each panel sub-title. The estimated trajectory and 95% confidence interval from the linear mixed models are plotted against years from baseline or symptom (MCI or AD) onset for each marker. Red line and pink shaded area represent the elevated amyloid subjects. Blue line and blue shaded area represent the normal amyloid subjects. L.R.= likelihood ratio

The CN, CN2MCI, and MCI2AD subgroups’ medians (interquartile range [IQR]) annual change in ADAS13 were (0.388 [−0.278, 0.818], 1.000 [0.239, 2.330], and 3.388 [1.750, 6.169], *p* < .001, respectively). The annual changes in CDRSB for each group were (0.000 [0.000, 0.000], 0.214 [0.100, 0.500], and 1.250 [0.750, 2.000], *p* < .001, respectively). The annual changes in MMSE for each group were (0.000 [−0.250, 0.161], −0.286 [−0.571, 0.000], and −1.500 [−2.775, −0.800], *p* < .001, respectively). The annual changes in hippocampal volume for each group were (−0.005 %ICV [−0.011, −0.001], −0.006 %ICV [−0.012, −0.002], and −0.014 %ICV [−0.021, −0.009], *p* < .001, respectively). The annual changes in FDG PET SUVR for each group were (−0.011 [−0.030, 0.010], −0.027 [−0.056, −0.012], and −0.039 [−0.063, 0.014], *p* < .001, respectively). The annual changes in Florbetapir PET SUVR for each group were (0.004 [−0.002, 0.012], 0.004 [−0.001,0.011], and 0.005 [−0.006, 0.014], *p* = .840, respectively). The annual changes in CSF Aβ_42_ for each group were (−1.500 pg/ml [−6.000, 4.000], −2.200 [−5.667, 4.000], and −2.000 [−7.000, 2.650], *p* = .564, respectively). The annual changes in CSF tau for each group were (0.775 pg/ml [−1.887, 4.500], 2.150 [−0.500, 7.900], and 3.000 [−3.900, 14.175], *p* = .121, respectively). The annual changes in CSF ptau for each group were (1.050 pg/ml [−1.450, 4.500], 1.980 [−0.200, 5.050], and 1.408 [−1.321, 8.325], *p* = .628, respectively).

Figure 2 shows the trajectories of the biomarker changes with either the normal or elevated amyloid groups. Qualitatively, the pattern remained stable in the CN, exhibited slow non-linear changes in the CN2MCI, and ended with a phase in which rapid non-linear changes appeared in the MCI2AD. We found no significant differences in the clinical profiles, hippocampal volume, or FDG changes between the elevated and normal amyloid subjects at the *p* < .05 level. The statistical results showed no difference for CDRSB and FDG in any of the three (CN, CN2MCI, and MCI2AD) subgroups at *p* < .05. The ADAS13 analysis showed significant group differences for the 6-9-year time period in the CN subgroup, for the < −4.5 and > 4 years to onset time in the CN2MCI subgroup, and for the > −0.5 years to onset time in the MCI2AD subgroup at *p* < .05. The MMSE analysis showed a significant group difference for the time period > −1 year in the CN2MCI subgroup at *p* < .05. Although the likelihood ratio test showed a significant difference in hippocampal volume between the elevated and normal amyloid subjects (*p* =.047), the post-hoc results showed no significance at *p* < .05 in the CN2MCI and only showed a significant group difference for the time period < −1 years in the MCI2AD at *p* < .05. All subgroups showed obvious significant differences with respect to florbetapir PET and CSF Aβ_42_ between the elevated and normal amyloid subjects at *p* < .001. For CSF Tau and CSF Ptau, only the CN2MCI subgroup showed no amyloid effect at *p* < .05; the other two subgroups showed significant differences at *p* < .05 (Figure 2; for the post hoc analysis results, see eTables 10-15).

We found that the changes in the CDRSB (Figure 3.1A and 2A), MMSE (Figure 3.1B and 3.2B), hippocampal volume (Figure 3.1C and 3.2C), and FDG PET in the post-cingulate cortex (Figure 3.1D and 3.2D) were associated with the change in the ADAS13 in both the CN2MCI and MCI2AD subgroups. However, the changes in the amyloid related biomarkers florbetapir PET and CSF Aβ_42_ were not significantly associated with the change in the ADAS13 in either group (Figure 3.1 E, 3.1F, 3.2 E, and 3.2 F).

**Figure 3.**
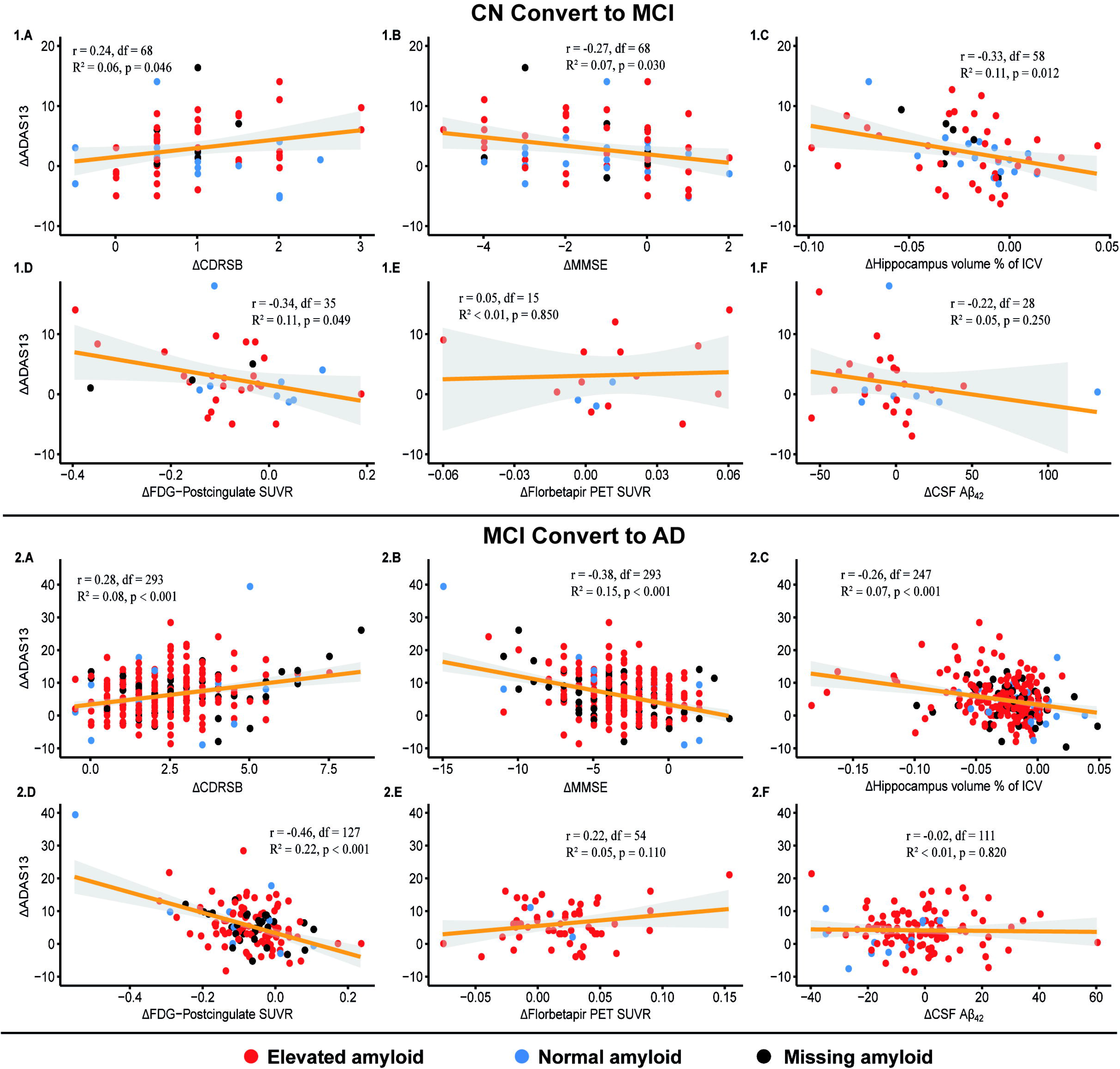
Relationship between the change in each biomarker and the change in ADAS13 in the CN conversion to MCI and the MCI conversion to AD subgroups. The top panels show that the changes in the (1.A) CDRSB score, (1.B) MMSE score, (1.C) hippocampal volume percent of ICV, and (1.D) post-cingulate FDG SUVR value significantly correlated with the change in the ADAS13 scores in the CN conversion to MCI subgroup. However, the changes in the amyloid-related biomarkers, (1.E) Florbetapir PET SUVR and (1.F) CSF Aβ_42_, were not significantly correlated with the change in ADAS13 scores. The bottom panels show that the change in the (2.A) CDRSB score, (2.B) MMSE score, (2.C) hippocampal volume percent of ICV, and (2.D) post-cingulate FDG SUVR value significantly correlated with the change in ADAS13 scores in the MCI conversion to AD subgroup. However, the changes in amyloid related biomarkers, (2.E) Florbetapir PET SUVR and (2.F) CSF Aβ_42_, were not significantly correlated with the change in ADAS13 scores. df = degree of freedom

Combining the biomarker findings, we assessed the trajectories and order of pathophysiological changes for the clinical, imaging, and biochemical measures (Figure 4.A and 4.B). As can be seen in Figures 1 and 2, the clinical profiles, hippocampal volume, and FDG changed slowly in the initial stage of CN2MCI and accelerated in the late MCI2AD stage. The order in which these measures changed in the CN2MCI subgroup was that the hippocampus and FDG PET changed earlier than ADAS13 and that CDRSB and MMSE were the last measures to change. Further, a *post hoc* analysis showed that the change in hippocampal volume preceded the symptom onset of MCI by 2.5 years and ADAS13 preceded the symptom onset of MCI by 1 year. Significant changes in MMSE and CDRSB were concurrent with MCI onset (Supplement Figure 3). Even in patients with elevated amyloid, the trajectory of the amyloid-related biomarker was not consistent with the clinical profiles, hippocampal volume, or FDG (Figure 4.B). More importantly, florbetapir PET was stable during the CN2MCI stage. Although CSF Aβ_42_ showed some nonlinear changes before MCI onset, the change was smaller than those of the other biomarkers. Thus, these results do not support previous reports^11,12^, suggesting that amyloid-related biomarker changes largely lead other biomarker changes at the onset of the disease.

**Figure 4.**
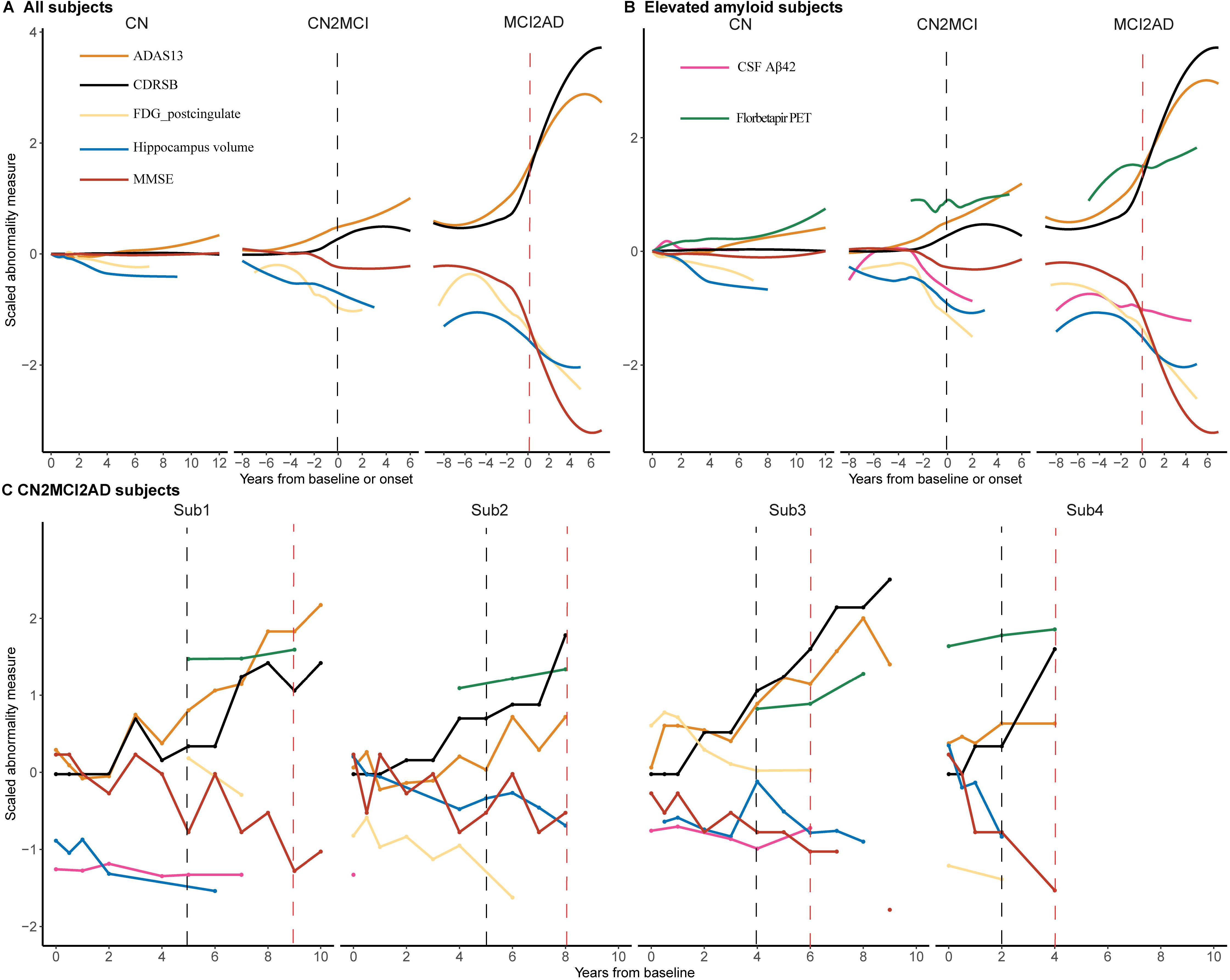
Temporal evolution of marker changes and within-individual trajectories of marker changes. Raw data for each biomarker and clinical profile converted to scaled values. The scaled value for each marker was defined by: (raw data – mean CN baseline value) / the standard deviation of the whole dataset. Clinical profiles, hippocampal volume, and FDG scaled changes in all subjects; Clinical profiles and biomarkers scaled changes in the elevated amyloid subjects; Clinical profiles and biomarkers scaled changes in 4 subjects who included the entire disease process from CN conversion to MCI followed by conversion to AD (CN2MCI2AD), within-individual changes.

We further assessed each biomarker for the individuals who progressed from CN to MCI and to AD for each biomarker (Figure 4.C and eFigure 4). The mean time for conversions from MCI to AD was 2.44±1.49 (range 1-7) years in these 24 subjects. The individual results were consistent with the previous group results: The trajectories of their clinical profiles changed slowly in the initial period in the CN2MCI stage and accelerated in the MCI2AD stage, the dynamic changes of hippocampal volume paralleled the disease status changes, and there were no significant changes in amyloid-related biomarkers in the CN to MCI to AD progression.

## Discussion

Identifying the dynamic changes in clinical assessments and biomarkers during a patient’s progression to AD is critical for defining the stage of the disease and its etiology and for monitoring the efficacy of potential therapies. In the present study, we avoided preconceptions about disease etiology and aligned the clinical symptom onset timepoints of the different stages from CN, through MCI, to AD using various clinical assessments and biomarkers to obtain a panorama of disease progression. The end stage of CN remained stable during the follow-up. Furthermore, the onset timepoints of clinical diagnosis were rigidly aligned for CN2MCI and MCI2AD with error less than 1 year. In addition, we also replicated the group results using the CN2MCI2AD subjects. Thus, the study expands the current literature about the SAD clinical and biomarker dynamic changes.

We found that the accumulation of amyloid in the CN did not predict future cognitive impairment in either people who maintained a stable CN or those in the pre-MCI onset stage (Figure 2.A-C). This result is consistent with recent reports that indicated that brain Aβ is not clinically relevant^28,29^. Other studies, however, reported that elevated amyloid in CN individuals was associated with a higher likelihood of cognitive decline compared with normal amyloid CN subjects^27,30^. Although these findings are insightful, using the same ADNI dataset, we found that cognitive decline did not depend on the accumulation of amyloid but on the clinical stage of the disease. Moreover, our results were partially consistent with previous findings, showing that amyloid accumulation alone cannot predict the cognitive decline or disease progression, whereas the amyloid accumulation combined with tau or atrophy of some groups can predict the further cognitive decline^31–33^ or disease progression in the normal adults^34–36^. These results support the recent idea about the combination therapies for future AD treatment strategies^37^ and the widespread concern about overdiagnosis in the preclinical AD^38^.

By aligning the disease onset timepoint, we found that dynamic changes in amyloid-related biomarkers were not associated with a change in disease status even in elevated amyloid subjects (Figures 2.F, 2.G, 4.B, 4.C). A previous prospective study, based on the amyloid hypothesis, reported that brain Aβ deposition continuously changed with SAD progression^18^. However, they found that the raw data of Aβ deposition was stable and changed slowly^18^, a finding that is in keeping with our results. In addition, by assessing the full range from CN to MCI to AD, we found that the trajectory of hippocampal volume and FDG were consistent with the clinical profiles in that they did not follow a sigmoid curve^20,21^ but rather showed a slow change in the initial stage and accelerated changes in the later stage from MCI to AD (Figure 4). Although previous studies based on the ADNI dataset reported that the changes in these biomarkers followed a sigmoid curve^39–41^, these studies did not align their findings with the stage of disease, so they could not be considered to accurately reflect the trajectory of biomarker changes that occur in the progression of AD. Thus, these results indicated that assessment of the clinical or biomarker dynamic changes by aligning the clinical onset timepoint maybe better reflect and characterize the progress of the disease.

Our finding that cognitive decline and Aβ deposition did not occur in parallel (Figures 3 and 4) is consistent with previous studies that reported that Aβ dysregulation poorly correlates with AD severity^42^, progressive neurodegeneration^43^, cognitive dysfunction^44^, or brain atrophy^45^. During the rapid cognitive decline from MCI to AD, Aβ deposition only mildly increased. This may partially explain why anti-Aβ drugs have failed in clinical trials. Medications, such as solanezumab, a medication designed to clear soluble Aβ from the brain, are used in the mild AD stage^8^, which is too late to prevent rapid cognitive decline. In addition, ADAS13 showed dramatic changes about 1 year before the clinical MCI onset, a finding which was not consistent with the general concept that clinical profiles change only after the onset of MCI^20,21^. Thus, the slow stage from pre-MCI to pre-AD may be a better time window for future clinical trial design.

Our results suggest that applying ADAD results directly to SAD research may not be appropriate^10^. We found that the rate of Aβ biomarker changes during CN conversion to MCI stage did not reflect those of other biomarkers and were not associated with clinical changes (Figure 4). This result is not consistent with previous ADAD studies that found that amyloid biomarkers undergo greater changes and lead to other biomarker changes in the initial stage of symptom onset^11–13^. The most likely explanation for this difference is that the ADAD and SAD have different etiologies^10^. In addition, we found that dramatic hippocampal atrophy starts 2.5 years prior to MCI onset, which is later than recent ADAD brain atrophy findings^13,14^. The concept that SAD involves a long pre-symptomatic period and is derived from ADAD studies^15^ may need to be reconsidered.

## Limitations

One of the limitations of the current study is that the CN2MCI subgroup was older than the MCI2AD subgroup, which may have influenced the pattern of biomarker changes. The ongoing ADNI dataset maybe resolve this limitation in future studies. Another limitation is the small sample size of the tau and Aβ biomarkers in the pre-MCI stage, which meant that we could not fully reveal the dynamic changes in these biomarkers in the preclinical stage. The ongoing collection of plasma biomarkers^46–50^ and ADNI3 tau-related PET data^51^ will improve the likelihood of fully understanding the preclinical stage of SAD in the future.

## Conclusions

Hippocampal volume and FDG associated with clinical profiles impairment in the SAD progression. Aβ alone is not associated with clinical profiles, hippocampal volume, and FDG impairment in the preclinical stage of SAD.

## Author Contributions

T.J. and P.F.B. supervised the study. J.Z., P.F.B., and T.J. were responsible for the design of the concept and the study. J.Z. contribution to the data analysis and statistical analysis; Y.Z., B.L., Y.L., and X.Z. made substantial contributions to the discussion on the results and the manuscript; J.Z., P.F.B., and T.J. wrote the manuscript.

## Acknowledgments

This work was partially supported by the Natural Science Foundation of China (Grant Nos. 31620103905 and 81701781), the Science Frontier Program of the Chinese Academy of Sciences (Grant No. QYZDJ-SSW-SMC019), the Guangdong Pearl River Talents Plan (2016ZT06S220), and the International Postdoctoral Exchange Fellowship Program 2017 by the Office of China Postdoctoral Council.

Data collection and sharing for this project was funded by the Alzheimer's Disease Neuroimaging Initiative (ADNI) (National Institutes of Health Grant U01 AG024904) and DOD ADNI (Department of Defense award number W81XWH-12-2-0012). ADNI is funded by the National Institute on Aging, the National Institute of Biomedical Imaging and Bioengineering, and through generous contributions from the following: AbbVie, Alzheimer’s Association; Alzheimer’s Drug Discovery Foundation; Araclon Biotech; BioClinica, Inc.; Biogen; Bristol-Myers Squibb Company; CereSpir, Inc.; Cogstate; Eisai Inc.; Elan Pharmaceuticals, Inc.; Eli Lilly and Company; EuroImmun; F. Hoffmann-La Roche Ltd and its affiliated company Genentech, Inc.; Fujirebio; GE Healthcare; IXICO Ltd.; Janssen Alzheimer Immunotherapy Research & Development, LLC.; Johnson & Johnson Pharmaceutical Research & Development LLC.; Lumosity; Lundbeck; Merck & Co., Inc.; Meso Scale Diagnostics, LLC.; NeuroRx Research; Neurotrack Technologies; Novartis Pharmaceuticals Corporation; Pfizer Inc.; Piramal Imaging; Servier; Takeda Pharmaceutical Company; and Transition Therapeutics. The Canadian Institutes of Health Research is providing funds to support ADNI clinical sites in Canada. Private sector contributions are facilitated by the Foundation for the National Institutes of Health (www.fnih.org). The grantee organization is the Northern California Institute for Research and Education, and the study is coordinated by the Alzheimer’s Therapeutic Research Institute at the University of Southern California. ADNI data are disseminated by the Laboratory for Neuro Imaging at the University of Southern California.

We thank Dr. Alan Ho who help statistical analysis, Dr. Wen Zhang who help access the data, Dr. Lingzhong Fan who offered constructive comments, and Dr. Rhoda E., Edmund F. Perozzi, and Rowan Tweedale for their editing assistance and discussions.

