## Supplement information for "Clinical and biomarker changes in sporadic Alzheimer’s disease"

### Supplementary methods

#### Details of the assessment of biomarkers and clinical profiles

All the clinical profiles and MRI-based volumetric data were extracted from the merged ADNI dataset (‘ADNIMERGE.csv’). The FDG data were extracted from the latest available dataset (‘UCBERKELEYFDG_09_05_18.csv’). The florbetapir PET data were extracted from the latest available dataset (‘UCBERKELEYAV45_04_12_19.csv’). Cerebrospinal fluid (CSF) data were obtained from the file ‘UPENNBIOMK_MASTER.csv’.

The clinical profiles and cognitive scores used in the present study included: the 13-item cognitive subscale of the Alzheimer’s Disease Assessment Scale (ADAS13), the Mini-Mental State Examination (MMSE), and the Clinical Dementia Rating Scale-Sum of Boxes (CDRSB). The structural MRI brain scans were acquired using 1.5 T and 3 T MRI scanners. Automated volume measures of MRI structure data were performed with FreeSurfer. In the present study, the sum of the left and right hippocampal volumes, adjusted for the total intracranial volume, was used to assess the degree of brain atrophy.

Fluorodeoxyglucose (FDG) positron emission tomography (PET) was used to measure cerebral metabolism. For the post-cingulate region, which is known to be an area of early deposition in ADAD[1,2], the standardized uptake value ratio (SUVR) was normalized with the entire brain stem as the reference region. To improve the accuracy of the longitudinal florbetapir PET change measurements, florbetapir PET SUVR was used as a weighted florbetapir mean in the frontal, cingulate, parietal, and temporal regions relative to the mean of the composite reference region (average of the whole cerebellum, brainstem/pons, and eroded white matter)[3]. The median value of all batches of the CSF tau, phosphor-tau (Ptau), and $A\beta_{42}$ were used in the present study.

### Supplementary figures

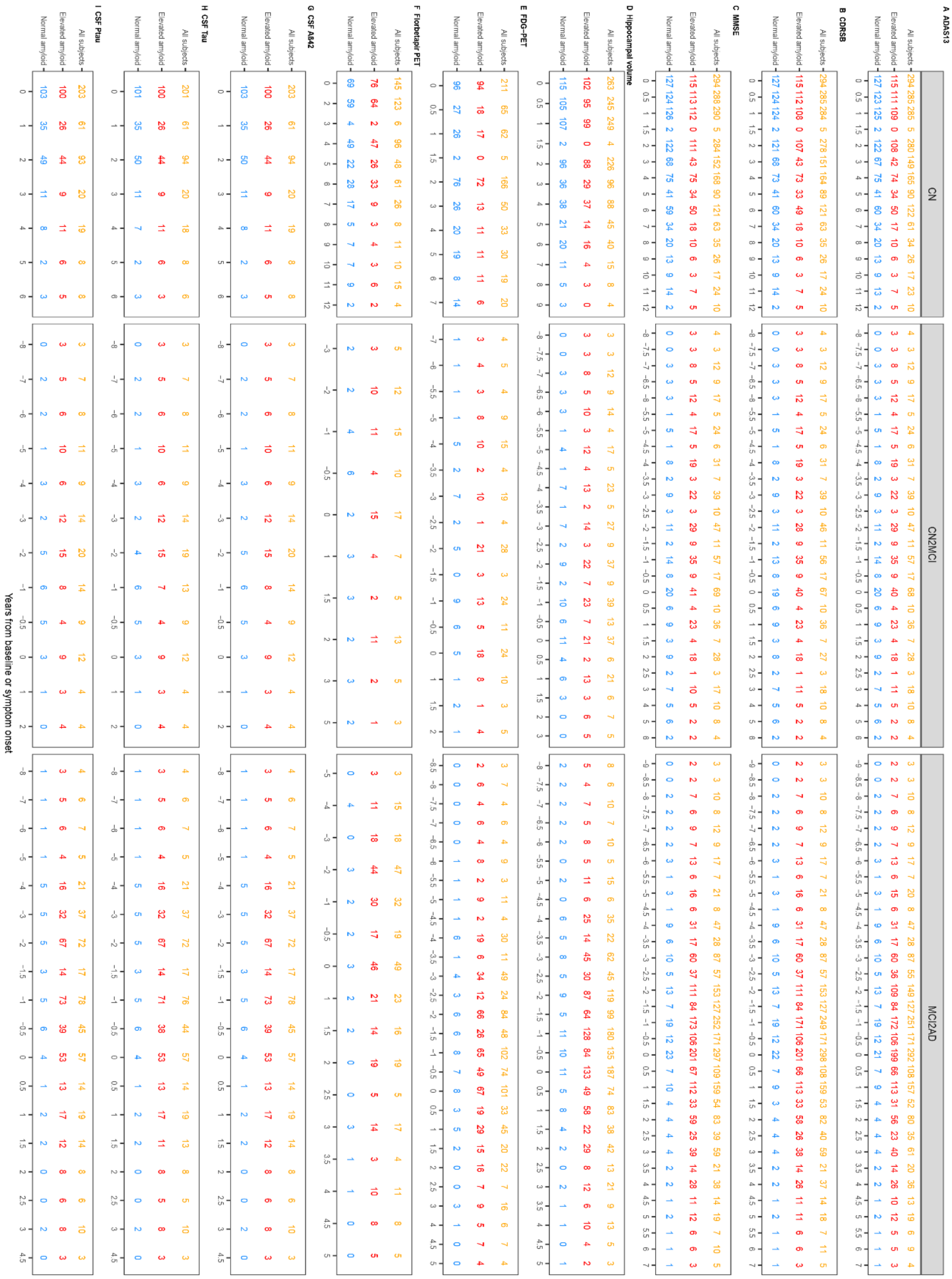

eFigure 1. Sample size at all timepoints for each biomarker used in the current study

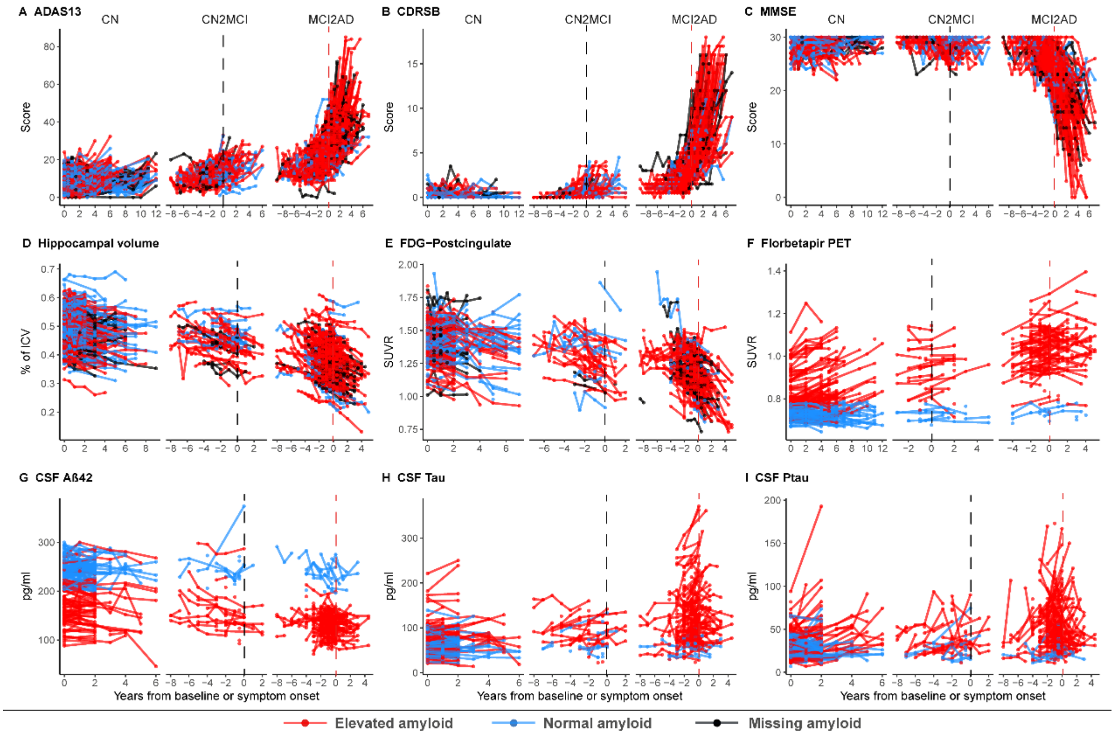

#### eFigure 2. Spaghetti plot of the raw data for each biomarker

See Figure 1 for an explanation of each panel subtitle.

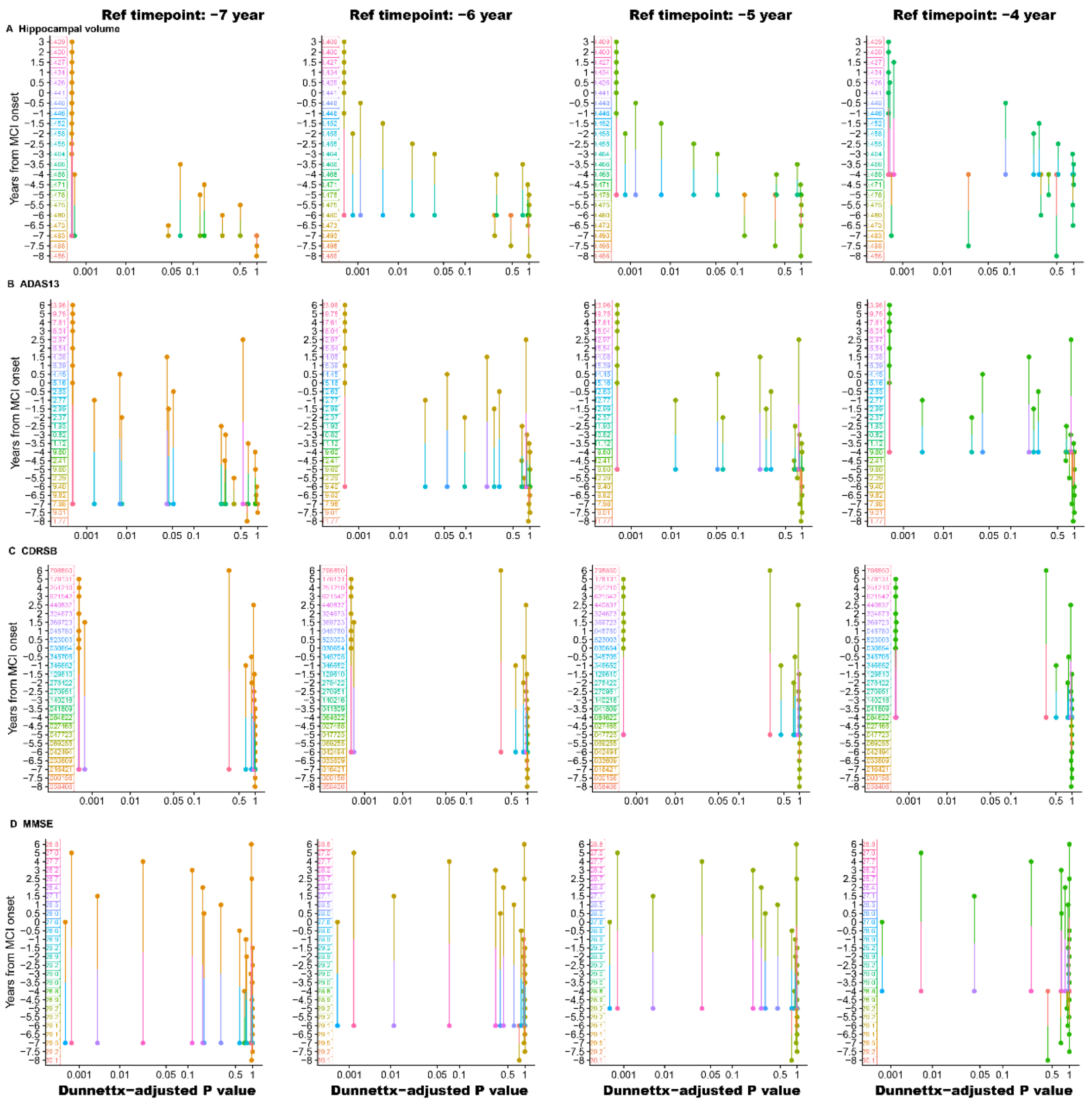

**eFigure 3. Post hoc analysis results between the pre-MCI onset timepoint and all the CN2MCI subgroup timepoints for the clinical profiles and hippocampal volume**

The x-axis of each panel is the Dunnettx-adjusted *p* value, and the y-axis is the years from MCI onset timepoint for hippocampal volume (Row A), ADAS13 (Row B), CDRSB (Row C), and MMSE (Row D). Each column represents the reference time for a stable stage timepoint in the CN2MCI subgroup. The line between two points in each panel indicates the *p* value of the *t* test for each marker between the two timepoints. The line located to the left of the *p* value on the x-axis indicates the significance level of the post hoc results between these two timepoints.

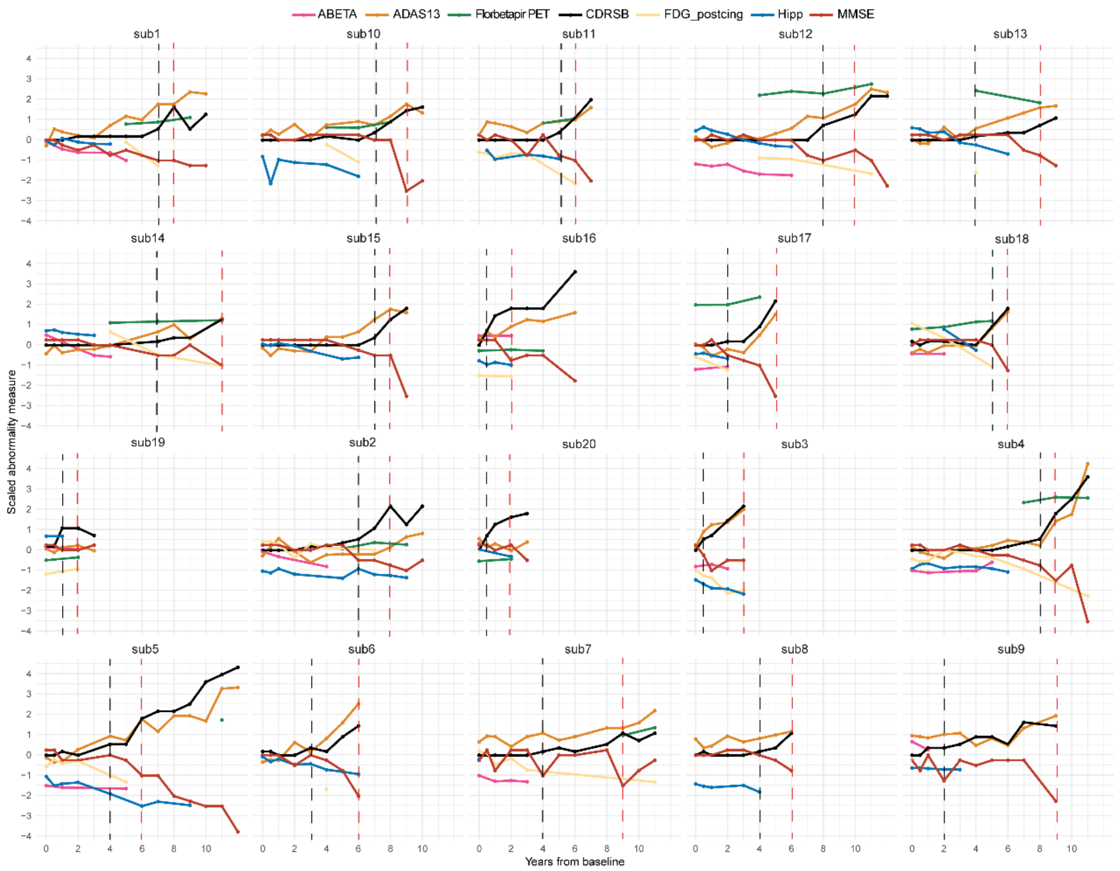

**eFigure 4. The within-individual trajectories for clinical and biomarkers in all CN2MCI2AD subjects**

The clinical profiles and biomarkers scaled changes for 20 subjects who had data that included all the disease stages from CN conversion to MCI and finally AD (CN2MCI2AD) within-individual changes. The results for the other 4 CN2MCI2AD individuals are shown in Figure 4C. The calculation of the scaled value is consistent with Figure 4.

### Supplementary tables

eTable 1 Linear Mixed-Effects Model of ADAS13

| Fixed effects | Coefficient (95% confidence limits) |
|---|---|
| Time | -0.554* (-1.000, -0.108) |
| $Time^2$ | 0.204*** (0.096, 0.311) |
| $Time^3$ | -0.009* (-0.016, -0.002) |
| Age (years) | 0.103*** (0.046, 0.160) |
| Education (years) | -0.185** (-0.326, -0.045) |
| CN2MCI | 5.145*** (3.625, 6.666) |
| MCI2AD | 18.301*** (17.359, 19.242) |
| Time:CN2MCI | 1.959*** (1.275, 2.643) |
| Time:MCI2AD | 4.771*** (4.253, 5.289) |
| $Time^2$:CN2MCI | -0.056 (-0.180, 0.067) |
| $Time^2$:MCI2AD | 0.048 (-0.066, 0.162) |
| $Time^3$:CN2MCI | 0.014* (0.001, 0.027) |
| $Time^3$:MCI2AD | -0.005 (-0.014, 0.005) |
| (Intercept) | 3.892 (-1.004, 8.787) |
| **Random effects** | **Standard deviation** |
| Time | 1.761 |
| $Time^2$ | 0.063 |
| $Time^3$ | 0.015 |
| (Intercept) | 5.132 |
| Residual | 3.156 |
| Number of individuals | 663 |
| Number of observations | 4,241 |
| Log likelihood | -12,476.120 |
| Akaike Inf. Crit. | 25,004.250 |
| Bayesian Inf. Crit. | 25,169.410 |
| Correlation Structure: Continuous AR(1) | |
| Note: | * *p* < .05; ** *p* < .01; *** *p* < .001 |

eTable 2 Linear Mixed-Effects Model of CDRSB

| Fixed effects | Coefficient (95% confidence limits) |
|---|---|
| Time | -0.013 (-0.118, 0.091) |
| $Time^2$ | 0.006 (-0.019, 0.032) |
| $Time^3$ | -0.0003 (-0.002, 0.001) |
| Male | -0.027 (-0.147, 0.094) |
| Education (years) | 0.006 (-0.016, 0.027) |
| CN2MCI | 0.780*** (0.517, 1.044) |
| MCI2AD | 3.933*** (3.758, 4.107) |
| APOE4 carriers | 0.034 (-0.093, 0.162) |
| Time:CN2MCI | 0.359*** (0.190, 0.528) |
| Time:MCI2AD | 1.397*** (1.273, 1.521) |
| $Time^2$:CN2MCI | 0.022 (-0.011, 0.055) |
| $Time^2$:MCI2AD | 0.138*** (0.110, 0.166) |
| $Time^3$:CN2MCI | -0.001 (-0.004, 0.003) |
| $Time^3$:MCI2AD | -0.001 (-0.004, 0.001) |
| (Intercept) | -0.017 (-0.391, 0.357) |
| **Random effects** | **Standard deviation** |
| Time | 0.462 |
| $Time^2$ | 0.049 |
| $Time^3$ | 0.003 |
| (Intercept) | 0.842 |
| Residual | 0.772 |
| Number of individuals | 663 |
| Number of observations | 4,241 |
| Log likelihood | -6,067.202 |
| Akaike Inf. Crit. | 12,188.400 |
| Bayesian Inf. Crit. | 12,360.060 |
| Correlation Structure: Continuous AR(1) | |

Note: *** $p < .001$

eTable 3 Linear Mixed-Effects Model of MMSE

| Fixed effects | Coefficient (95% confidence limits) |
|---|---|
| Time | 0.045 (-0.150, 0.239) |
| $Time^2$ | -0.021 (-0.070, 0.027) |
| $Time^3$ | 0.001 (-0.002, 0.004) |
| Age (years) | -0.017 (-0.035, 0.001) |
| Education (years) | 0.095*** (0.051, 0.138) |
| CN2MCI | -0.647** (-1.132, -0.163) |
| MCI2AD | -4.914*** (-5.238, -4.590) |
| APOE4 carriers | 0.149 (-0.110, 0.408) |
| Time:CN2MCI | -0.331* (-0.619, -0.043) |
| Time:MCI2AD | -1.788*** (-2.011, -1.565) |
| $Time^2$:CN2MCI | -0.001 (-0.059, 0.058) |
| $Time^2$:MCI2AD | -0.121*** (-0.173, -0.070) |
| $Time^3$:CN2MCI | -0.002 (-0.008, 0.004) |
| $Time^3$:MCI2AD | 0.004 (-0.0002, 0.008) |
| (Intercept) | 28.724*** (27.184, 30.264) |
| **Random effects** | **Standard deviation** |
| Time | 0.687 |
| $Time^2$ | 0.054 |
| $Time^3$ | 0.006 |
| (Intercept) | 1.454 |
| Residual | 1.578 |
| Number of individuals | 663 |
| Number of observations | 4,292 |
| Log likelihood | -8,964.928 |
| Akaike Inf. Crit. | 17,983.860 |
| Bayesian Inf. Crit. | 18,155.700 |
| Correlation Structure: Continuous AR(1) | |
| Note: | * $p < .05$; ** $p < .01$; *** $p < .001$ |

eTable 4 Linear Mixed-Effects Model of Hippocampal volume

| Fixed effects | Coefficient (95% confidence limits) |
|---|---|
| Time | -0.005*** (-0.006, -0.004) |
| $Time^3$ | 0.00004** (0.00002, 0.0001) |
| CN2MCI | -0.038*** (-0.053, -0.023) |
| MCI2AD | -0.116*** (-0.126, -0.106) |
| 3T MRI | 0.024*** (0.019, 0.029) |
| Age (years) | -0.004*** (-0.004, -0.003) |
| Male | -0.017*** (-0.026, -0.009) |
| Education (years) | -0.003*** (-0.005, -0.002) |
| APOE4 carriers | -0.008 (-0.017, 0.001) |
| Time:CN2MCI | -0.004** (-0.006, -0.001) |
| Time:MCI2AD | -0.010*** (-0.011, -0.008) |
| Time:3T MRI | -0.002** (-0.003, -0.001) |
| $Time^3$:3T MRI | -0.00004* (-0.0001, -0.00000) |
| (Intercept) | 0.816*** (0.762, 0.871) |
| **Random effects** | **Standard deviation** |
| Time | 0.005 |
| $Time^2$ | <0.001 |
| (Intercept) | 0.053 |
| Residual | 0.012 |
| Number of individuals | 646 |
| Number of observations | 2,846 |
| Log likelihood | 6,967.814 |
| Akaike Inf. Crit. | -13,893.630 |
| Bayesian Inf. Crit. | -13,768.600 |

Note: * $p < .05$; ** $p < .01$; *** $p < .001$

eTable 5 Linear Mixed-Effects Model of FDG

| Fixed effects | Coefficient (95% confidence limits) |
|---|---|
| Time | -0.005 (-0.011, 0.001) |
| $Time^2$ | -0.002*** (-0.002, -0.001) |
| $Time^3$ | 0.0001* (0.00002, 0.0002) |
| Age (years) | -0.004*** (-0.006, -0.002) |
| Male | 0.018 (-0.006, 0.042) |
| CN2MCI | -0.133*** (-0.175, -0.091) |
| MCI2AD | -0.231*** (-0.260, -0.202) |
| APOE4 carriers | -0.024 (-0.050, 0.003) |
| Time:CN2MCI | -0.029*** (-0.041, -0.018) |
| Time:MCI2AD | -0.039*** (-0.047, -0.031) |
| (Intercept) | 1.719*** (1.584, 1.854) |
| **Random effects** | **Standard deviation** |
| Time | 0.017 |
| (Intercept) | 0.127 |
| Residual | 0.054 |
| Number of individuals | 522 |
| Number of observations | 1,567 |
| Log likelihood | 1,595.474 |
| Akaike Inf. Crit. | -3,160.947 |
| Bayesian Inf. Crit. | -3,080.593 |
| Correlation Structure: Continuous AR(1) | |

Note: * $p < .05$; *** $p < .001$

eTable 6 Linear Mixed-Effects Model of Florbetapir PET

| Fixed effects | Coefficient (95% confidence limits) |
| --- | --- |
| Time | 0.009*** (0.005, 0.013) |
| Time$^2$ | -0.0003 (-0.001, 0.001) |
| Time$^3$ | 0.0001 (-0.00001, 0.0001) |
| Normal amyloid | -0.093*** (-0.117, -0.069) |
| CN2MCI | 0.116*** (0.084, 0.148) |
| MCI2AD | 0.204*** (0.181, 0.227) |
| Male | -0.013 (-0.030, 0.004) |
| Education (years) | 0.003 (-0.0003, 0.006) |
| APOE4 carriers | 0.022* (0.002, 0.041) |
| Time:Normal Amyloid | -0.008*** (-0.013, -0.004) |
| Time$^2$:Normal Amyloid | -0.0001 (-0.001, 0.001) |
| Time$^3$:Normal Amyloid | -0.00003 (-0.0001, 0.00004) |
| Time:CN2MCI | -0.002 (-0.009, 0.004) |
| Time:MCI2AD | -0.005* (-0.010, -0.001) |
| Time$^2$:CN2MCI | -0.0002 (-0.002, 0.001) |
| Time$^2$:MCI2AD | -0.0004 (-0.001, 0.001) |
| Time$^3$:CN2MCI | -0.00000 (-0.001, 0.001) |
| Time$^3$:MCI2AD | 0.0001 (-0.0001, 0.0002) |
| Normal amyloid:CN2MCI | -0.119*** (-0.174, -0.063) |
| Normal amyloid:MCI2AD | -0.193*** (-0.245, -0.142) |
| Time$^3$:Normal amyloid:CN2MCI | 0.0001 (-0.001, 0.001) |
| Time$^3$:Normal amyloid:MCI2AD | 0.001 (-0.0002, 0.001) |
| (Intercept) | 0.780*** (0.726, 0.835) |
| **Random effects** | **Standard deviation** |
| Time | 0.008 |
| Time$^3$ | <0.0001 |
| (Intercept) | 0.080 |
| Residual | 0.014 |
| Number of individuals | 397 |
| Number of observations | 936 |
| Log likelihood | 1,611.051 |
| Akaike Inf. Crit. | -3,160.101 |
| Bayesian Inf. Crit. | -3,010.011 |

Correlation Structure: Continuous AR(1)

Note: * *p* < .05; *** *p* < .001

eTable 7 Linear Mixed-Effects Model of CSF Aβ42

| Fixed effects | Coefficient (95% confidence limits) |
|---|---|
| Time | 1.408 (-1.803, 4.619) |
| Time$^2$ | -1.510*** (-2.338, -0.682) |
| Time$^3$ | 0.048 (-0.017, 0.114) |
| Normal amyloid | 68.215*** (60.032, 76.399) |
| CN2MCI | -19.422** (-32.480, -6.364) |
| MCI2AD | -34.859*** (-42.628, -27.090) |
| Age (years) | -0.608** (-1.009, -0.208) |
| APOE4 carriers | -12.349*** (-18.560, -6.138) |
| Time:Normal amyloid | -1.537 (-5.109, 2.036) |
| Time$^2$:Normal amyloid | 0.851* (0.173, 1.529) |
| Time:CN2MCI | -3.576 (-8.590, 1.437) |
| Time:MCI2AD | -3.012 (-6.755, 0.731) |
| Time$^2$:CN2MCI | 2.154** (0.800, 3.507) |
| Time$^2$:MCI2AD | 1.649** (0.576, 2.723) |
| Normal amyloid:CN2MCI | 34.087** (12.736, 55.438) |
| Normal amyloid:MCI2AD | 14.545 (-2.598, 31.689) |
| Time:Normal amyloid:CN2MCI | 14.904*** (6.693, 23.115) |
| Time:Normal amyloid:MCI2AD | -1.403 (-7.782, 4.976) |
| (Intercept) | 221.314*** (190.706, 251.922) |
| **Random effects** | **Standard deviation** |
| Time | 3.512 |
| (Intercept) | 26.673 |
| Residual | 12.181 |
| Number of individuals | 451 |
| Number of observations | 951 |
| Log likelihood | -4,285.484 |
| Akaike Inf. Crit. | 8,616.967 |
| Bayesian Inf. Crit. | 8,728.690 |

Note: * *p* < .05; ** *p* < .01; *** *p* < .001

eTable 8 Linear Mixed-Effects Model of CSF PTAU

| Fixed effects | Coefficient (95% confidence limits) |
|---|---|
| Time | 2.174 (-2.474, 6.822) |
| Time$^2$ | 1.045 (-1.264, 3.354) |
| Time$^3$ | -0.097 (-0.390, 0.197) |
| Normal amyloid | -5.911 (-11.859, 0.037) |
| CN2MCI | 14.572** (4.891, 24.253) |
| MCI2AD | 25.201*** (19.482, 30.919) |
| Male | -3.072 (-6.985, 0.841) |
| Education (years) | 0.365 (-0.326, 1.056) |
| APOE4 carriers | 0.064 (-4.222, 4.349) |
| Time:Normal amyloid | -3.232 (-6.911, 0.448) |
| Time$^2$:Normal amyloid | 0.024 (-1.132, 1.180) |
| Time$^3$:Normal amyloid | -0.013 (-0.188, 0.162) |
| Time:CN2MCI | -0.888 (-7.024, 5.248) |
| Time:MCI2AD | -0.147 (-5.030, 4.736) |
| Time$^2$:CN2MCI | -1.169 (-3.730, 1.392) |
| Time$^2$:MCI2AD | -1.104 (-3.417, 1.209) |
| Time$^3$:CN2MCI | 0.096 (-0.223, 0.415) |
| Time$^3$:MCI2AD | 0.107 (-0.194, 0.408) |
| Normal amyloid:CN2MCI | -9.381 (-25.708, 6.945) |
| Normal amyloid:MCI2AD | -25.256*** (-38.264, -12.248) |
| Time:Normal amyloid:CN2MCI | 3.485 (-5.144, 12.114) |
| Time:Normal amyloid:MCI2AD | 2.736 (-3.426, 8.899) |
| Time$^3$:Normal amyloid:CN2MCI | 0.006 (-0.300, 0.313) |
| Time$^3$:Normal amyloid:MCI2AD | -0.022 (-0.324, 0.280) |
| (Intercept) | 27.688*** (15.783, 39.592) |
| **Random effects** | **Standard deviation** |
| Time | 5.143 |
| Time$^2$ | 0.080 |
| Time$^3$ | 0.078 |
| (Intercept) | 19.489 |
| Residual | 8.789 |
| Number of individuals | 451 |
| Number of observations | 950 |
| Log likelihood | -4,053.149 |
| Akaike Inf. Crit. | 8,180.297 |
| Bayesian Inf. Crit. | 8,359.986 |

Correlation Structure: Continuous AR(1)

| | |
|---|---|
| Note: | ** $p < .01$; *** $p < .001$ |

eTable 9 Linear Mixed-Effects Model of CSF TAU

| Fixed effects | Coefficient (95% confidence limits) |
|---|---|
| Time | 4.089*** (2.649, 5.529) |
| $Time^2$ | -0.378** (-0.634, -0.121) |
| $Time^3$ | -0.064** (-0.112, -0.017) |
| Normal amyloid | -7.950 (-20.622, 4.723) |
| CN2MCI | 41.014*** (23.599, 58.429) |
| MCI2AD | 64.650*** (53.384, 75.916) |
| Male | -12.525** (-20.750, -4.300) |
| Time:Normal amyloid | -4.256** (-7.054, -1.458) |
| $Time^2$:Normal amyloid | 0.484 (-0.043, 1.010) |
| $Time^3$:Normal amyloid | 0.097 (-0.002, 0.196) |
| Normal Amyloid:CN2MCI | -29.576* (-58.929, -0.224) |
| Normal Amyloid:MCI2AD | -63.073*** (-87.464, -38.682) |
| (Intercept) | 73.724*** (64.113, 83.336) |
| **Random effects** | **Standard deviation** |
| Time | 6.105 |
| $Time^2$ | 0.764 |
| $Time^3$ | 0.142 |
| (Intercept) | 44.415 |
| Residual | 11.171 |
| Number of individuals | 446 |
| Number of observations | 939 |
| Log likelihood | -4,413.427 |
| Akaike Inf. Crit. | 8,874.855 |
| Bayesian Inf. Crit. | 8,991.130 |
| Note: | * $p < .05$; ** $p < .01$; *** $p < .001$ |

**eTable 10.** *Post hoc* analysis of the relationship between the elevated amyloid and normal amyloid groups with respect to ADAS13 at each CN subgroup timepoint

| Timepoint | 0 | 0.5 | 1 | 2 | 3 | 4 | 5 | 6 | 7 | 8 | 9 | 10 | 11 | 12 |
|---|---|---|---|---|---|---|---|---|---|---|---|---|---|---|
| t.ratio | -0.873 | -0.818 | -0.662 | -0.099 | 0.640 | 1.319 | 1.810 | 2.116 | 2.270 | 2.276 | 2.097 | 1.668 | 0.985 | 0.200 |
| *p* | 0.384 | 0.414 | 0.509 | 0.921 | 0.523 | 0.189 | 0.072 | 0.035 | 0.024 | 0.024 | 0.037 | 0.097 | 0.326 | 0.842 |

**Supplementary Table 11.** *Post hoc* analysis of the relationship between the elevated amyloid and normal amyloid groups with respect to ADAS13 at each CN2MCI subgroup timepoint

| Timepoint | -7 | -6.5 | -6 | -5.5 | -5 | -4.5 | -4 | -3.5 | -3 | -2.5 | -2 | -1.5 |
|---|---|---|---|---|---|---|---|---|---|---|---|---|
| t.ratio | -2.794 | -2.676 | -2.523 | -2.327 | -2.084 | -1.794 | -1.463 | -1.103 | -0.730 | -0.364 | -0.019 | 0.296 |
| *p* | 0.007 | 0.010 | 0.014 | 0.024 | 0.042 | 0.078 | 0.149 | 0.275 | 0.468 | 0.717 | 0.985 | 0.769 |
| Timepoint | -1 | -0.5 | 0 | 0.5 | 1 | 1.5 | 2 | 3 | 4 | 5 | 6 | |
| t.ratio | 0.575 | 0.819 | 1.030 | 1.211 | 1.367 | 1.501 | 1.617 | 1.804 | 1.947 | 2.059 | 2.148 | |
| *p* | 0.568 | 0.416 | 0.307 | 0.231 | 0.177 | 1.389 | 0.111 | 0.076 | 0.056 | 0.044 | 0.036 | |

**eTable 12.** *Post hoc* analysis of the relationship between the elevated amyloid and normal amyloid groups with respect to ADAS13 at each MCI2AD subgroup timepoint

| Timepoint | -8 | -7.5 | -7 | -6.5 | -6 | -5.5 | -5 | -4.5 | -4 | -3.5 |
|---|---|---|---|---|---|---|---|---|---|---|
| t.ratio | -0.361 | 0.157 | 0.706 | 1.177 | 1.489 | 1.642 | 1.676 | 1.630 | 1.531 | 1.400 |
| *p* | 0.719 | 0.875 | 0.481 | 0.241 | 0.138 | 0.102 | 0.095 | 0.105 | 0.127 | 0.163 |
| Timepoint | -3 | -2.5 | -2 | -1.5 | -1 | -0.5 | 0 | 0.500 | 1.000 | 1.500 |
| t.ratio | 1.259 | 1.136 | 1.069 | 1.092 | 1.229 | 1.479 | 1.816 | 2.197 | 2.569 | 2.890 |
| *p* | 0.209 | 0.257 | 0.287 | 0.276 | 0.220 | 0.141 | 0.071 | 0.029 | 0.011 | 0.004 |
| Timepoint | 2.000 | 2.500 | 3.000 | 3.500 | 4.000 | 4.500 | 5.000 | 5.500 | 6.000 | 7.000 |
| t.ratio | 3.140 | 3.317 | 3.433 | 3.504 | 3.543 | 3.560 | 3.563 | 3.556 | 3.545 | 3.513 |
| *p* | 0.002 | 0.001 | <0.001 | <0.001 | <0.001 | <0.001 | <0.001 | <0.001 | <0.001 | <0.001 |

**eTable 13.** *Post hoc* analysis of the relationship between the elevated amyloid and normal amyloid groups with respect to MMSE at each CN2MCI subgroup timepoint

| Timepoint | -7 | -6.5 | -6 | -5.5 | -5 | -4.5 | -4 | -3.5 | -3 | -2.5 | -2 | -1.5 |
|---|---|---|---|---|---|---|---|---|---|---|---|---|
| t.ratio | -1.367 | -1.333 | -1.252 | -1.105 | -0.933 | -0.816 | -0.793 | -0.867 | -1.029 | -1.269 | -1.574 | -1.922 |
| *p* | 0.177 | 0.188 | 0.216 | 0.274 | 0.355 | 0.418 | 0.431 | 0.389 | 0.308 | 0.210 | 0.121 | 0.060 |
| Timepoint | -1 | -0.5 | 0 | 0.5 | 1 | 1.5 | 2 | 2.5 | 3 | 4 | 5 | 6 |
| t.ratio | -2.279 | -2.604 | -2.865 | -3.049 | -3.163 | -3.225 | -3.251 | -3.255 | -3.240 | -3.149 | -2.890 | -2.297 |
| *p* | 0.026 | 0.012 | 0.006 | 0.004 | 0.003 | 0.002 | 0.002 | 0.002 | 0.002 | 0.003 | 0.005 | 0.025 |

**eTable 14.** *Post hoc* analysis of the relationship between the elevated amyloid and normal amyloid groups with respect to hippocampal volume at each CN2MCI subgroup timepoint

| Timepoint | -7 | -6.5 | -6 | -5.5 | -5 | -4.5 | -4 | -3.5 | -3 |
|---|---|---|---|---|---|---|---|---|---|
| t.ratio | 0.116 | 0.012 | -0.098 | -0.214 | -0.335 | -0.461 | -0.591 | -0.724 | -0.859 |
| *p* | 0.908 | 0.991 | 0.922 | 0.831 | 0.739 | 0.647 | 0.557 | 0.472 | 0.394 |
| Timepoint | -2.5 | -2 | -1.5 | -1 | -0.5 | 0 | 0.5 | 1 | 1.5 |
| t.ratio | -0.994 | -1.128 | -1.260 | -1.388 | -1.511 | -1.626 | -1.735 | -1.835 | -1.926 |
| *p* | 0.325 | 0.264 | 0.213 | 0.171 | 0.137 | 0.110 | 0.088 | 0.072 | 0.059 |

**eTable 15.** *Post hoc* analysis of the relationship between the elevated amyloid and normal amyloid groups with respect to hippocampal volume at each MCI2AD subgroup timepoint

| Timepoint | -8 | -7.5 | -7 | -6.5 | -6 | -5 | -4 | -3.5 | -3 | -2.5 | -2 | -1.5 |
|---|---|---|---|---|---|---|---|---|---|---|---|---|
| t.ratio | 4.412 | 4.364 | 4.300 | 4.217 | 4.114 | 3.836 | 3.452 | 3.220 | 2.965 | 2.691 | 2.403 | 2.107 |
| *p* | <0.001 | <0.001 | <0.001 | <0.001 | <0.001 | <0.001 | <0.001 | 0.002 | 0.003 | 0.008 | 0.017 | 0.036 |
| Timepoint | -1 | -0.5 | 0 | 0.5 | 1 | 1.5 | 2 | 3 | 3.5 | 4 | 5 | |
| t.ratio | 1.808 | 1.512 | 1.223 | 0.946 | 0.682 | 0.434 | 0.202 | -0.214 | -0.398 | -0.568 | -0.869 | |

| | | | | | | | | | | | |
|---|---|---|---|---|---|---|---|---|---|---|---|
| *p* | 0.072 | 0.132 | 0.223 | 0.345 | 0.496 | 0.665 | 0.840 | 0.831 | 0.691 | 0.571 | 0.386 |
